## Supplemental material for "Maximum *a posteriori* natural scene reconstruction from retinal ganglion cells with deep denoiser priors"

### A Appendix

#### A.1 Potential negative impacts

A potential negative impact of any animal research is use of the animals. In the present work, eyes were obtained from animals that were euthanized in the course of experiments already being performed by other laboratories. If the eyes had not been used for these recordings, they would have been discarded. We are unaware of any other significant potential negative impacts.

#### A.2 Summary of experimental datasets

Table 4: Size (number of stimuli) of dataset partitions

| | Train | Test | Heldout |
|---|---|---|---|
| Retina 1 | 17,500 | 1000 | 500 |
| Retina 2 | 9000 | 500 | 500 |

Table 5: Cell counts for each dataset

| | Total RGC count | ON parasol | OFF parasol | ON midget | OFF midget |
|---|---|---|---|---|---|
| Retina 1 | 695 | 79 | 115 | 228 | 273 |
| Retina 2 | 778 | 97 | 130 | 207 | 344 |

#### A.3 Computational resources

All computations were performed using Pytorch [52] on a single NVIDIA V100 GPU with 32 GB of VRAM. Fitting encoding models for all cells in a single retina required approximately 24 hours of computational time.

#### A.4 Linear-nonlinear-Poisson (LNP) model and fitting

The LNP response model for a single RGC is

$$p(s|\mathbf{x}) \sim \text{Poiss}\left(\exp(\mathbf{m}^T\mathbf{x} + b)\right). \tag{9}$$

To fit this model to spike count data, we optimize the parameters $\mathbf{m}$ (the spatial stimulus filter) and $b$ (the scalar bias) according to:

$$\arg\min_{\mathbf{m},b} \left\{ \exp\{\mathbf{m}^T\mathbf{x} + b\} - s(\mathbf{m}^T\mathbf{x} + b) + \gamma_1|\mathbf{m}|_1 + \frac{\gamma_2}{2}|\mathbf{m} - \mathbf{m}'|_2^2 \right\}. \tag{10}$$

The first two terms correspond to the negative log-likelihood of the model of equation (9). The third term is an L1 sparsity-inducing penalty on the spatial filter $\mathbf{m}$. The fourth term is an L2 penalty that induces similarity to $\mathbf{m}'$, which is an initial fit of the spatial filter obtained using reverse correlation with white noise. Together, the two regularization terms help to ensure that the spatial filters are spatially compact, contiguous, and resemble the white noise receptive fields. The hyperparameters for this objective function were optimized by performing a grid search and evaluating the log-likelihood of the responses of a small number of RGCs of each type in test partition. Optimization problems were solved separately for each RGC using FISTA [26].

#### A.5 Generalized linear model (GLM) and fitting

The GLM model of RGCs augments the LNP model to include feedback and coupling filters to account for refractoriness, bursting, and cell-to-cell correlations [10]. We make the following definitions and assumptions:

- The spikes for each RGC are binned into $T+1$ total time bins indexed as $t \in [0, 1, \ldots T-1, T]$ The stimulus onset (transition from gray screen to presented image) occurs at time $t = N$.
- The time bin durations are sufficiently brief (1 ms) that at most one spike per RGC can occur in any bin.
- The spike train for the $i^{\text{th}}$ single RGC is denoted $s_i$. The spike trains for a set of RGCs $A$ is denoted $\mathbf{s}_A$, and the spike trains for the set of all RGCs except cell $i$ is referred to as $\mathbf{s}_{\{i\}^C}$
- The visual stimulus within each trial is space-time separable, because each trial consists of a single statically-flashed image. In particular, the visual stimulus $v[x, y, t] = w[t]\mathbf{x}$ where $\mathbf{x}$ is the stimulus image, and $w[t]$ is a boxcar corresponding to the time component of the stimulus.

##### A.5.1 GLM with Bernoulli spiking

We assume that the spatio-temporal stimulus filter for the RGC is space-time separable. The GLM for the $i$th cell thus the following parameters:

- The spatial component of the stimulus filter $\mathbf{m}_i$
- The temporal component of the stimulus filter $h_i[t]$, assumed causal
- The additive (bias) constant $b_i$
- A feedback (spike history) filter $f_i[t]$, assumed causal
- Coupling filters for the neighboring RGCs, $c_i^{(j)}[t]$, assumed causal. The set of cells coupled to cell $i$ is denoted $\{i\}^C$

Approximate nearest-neighbor distances for each of the cell types (ON parasol, OFF parasol, ON midget, OFF midget) were computed based on the reverse-correlation receptive field centers. Coupled neighbors consisted of cells within a multiple of the median nearest neighbor distances for each respective cell type (2 times the median nearest neighbor distance for parasols, 2.5 times the median nearest neighbor distance for midgets).

The generator signal of the $i$th RGC, which determines its instantaneous spiking probability, is defined as

$$g_i[t] = (\mathbf{m}_i^T\mathbf{x})(h_i * w)[t-1] + (s_i * f_i)[t-1] + \sum_{j \in \{i\}^C} (s_j * c_i^{(j)})[t-1] + b_i. \tag{11}$$

Since filters $h_i$, $f_i$, and $c_i^{(j)}$ are causal, $g_i[t]$ depends only on the stimulus and spikes that occur before time $t$.

The generator signal is passed through a sigmoidal nonlinearity $p[t] = \exp\{g_i[t]\}/(1 + \exp\{g_i[t]\})$, and a spike is generated with this probability. The log-likelihood of observing $s_0[t]$ spikes from cell $i$ at time $t$ given the stimulus and previously observed spikes is

$$\log p(s_i[t] \mid \mathbf{x}, s_i[0, \ldots, t-1], \mathbf{s}_{\{i\}^C}[0, \ldots, t-1]) = s_i[t]g_i[t] - \log(1 + \exp\{g_i[t]\}) \tag{12}$$

##### A.5.2 Derivation of GLM log-likelihood objective function (equation (5))

When fitting optimal GLM parameters for the $i^{\text{th}}$ RGC, the spikes from the coupled RGCs $\mathbf{s}_{\{i\}^C}[0, \ldots, T]$ are observed and known, and the spikes for RGC $i$ that occur before time $N$ $s_i[0, \ldots, N-1]$ are also known. The optimal GLM parameters for the $i^{\text{th}}$ RGC $\mathbf{m}_i, h_i, f_i, c_i^{(j)}$, and $b_i$ can be found by minimizing the negative log-likelihood (negative of equation (12)) over the parameters. The negative log-likelihood can be simplified using the chain rule:

$$
\begin{aligned}
&-\log p(s_i[N,\ldots,T] \mid \mathbf{x}, s_i[0,\ldots N-1], \mathbf{s}_{\{i\}^C}[0,\ldots,T]) \\
&= -\sum_{t=N}^{T} \log p(s_i[t] \mid \mathbf{x}, s_i[0,\ldots,t-1], \mathbf{s}_{\{i\}^C}[0,\ldots,t-1]) \\
&= \sum_{t=N}^{T} \big\{ \log(1+\exp\{g_i[t]\}) - s_i[t] g_i[t] \big\},
\end{aligned}
$$

which results in the expression presented in (5).

##### A.5.3 Regularized GLM objective function

To reduce the number of GLM model parameters, the stimulus time filter $h_i[t]$, feedback (spike history) filter $f_i[t]$, and the neighboring cell coupling filters $c_i^{(j)}[t]$ were represented in terms of a small number of cosine bump basis vectors [10] (basis functions discussed in more detail in A.5.4). Letting $B_h$, $B_f$, and $B_c$ represent the basis matrices for stimulus time, feedback, and coupling filters, respectively, the full time-domain filters can be recovered from the low dimensional representations $\mathbf{h}_i = B_h \tilde{\mathbf{h}}_i$, $\mathbf{f}_i = B_h \tilde{\mathbf{f}}_i$, and $\mathbf{c}_i^{(j)} = B_c \tilde{\mathbf{c}}_i^{(j)}$.

Substituting these low-dimensional temporal filters into the encoding negative log-likelihood from equation (5) yields an expression for that is jointly convex in $\{\mathbf{m}_i, \tilde{\mathbf{f}}_i, \tilde{\mathbf{c}}_i^{(j)}, b_i\}$ and in $\{\tilde{\mathbf{h}}_i, \tilde{\mathbf{f}}_i, \tilde{\mathbf{c}}_i^{(j)}, b_i\}$ but not with the stimulus spatial filter $\mathbf{m}$ and stimulus temporal filter $\tilde{\mathbf{h}}_i$ together because of the space-time stimulus filter separability assumption. As with the LNP models, L1 sparsity and L2 prior regularization terms were added. In addition, to eliminate spurious cell-cell correlations, an $L_{2,1}$ group-sparsity term was added to constrain the neighboring cell coupling filters $\tilde{\mathbf{c}}_i^{(j)}, b_i$. Because of the joint convexity issue described above, we fitted GLMs by alternating between solving two convex minimization problems:

$$
\begin{aligned}
\arg \min_{\mathbf{m}_i, \tilde{\mathbf{f}}_i, \tilde{\mathbf{c}}_i^{(j)}, b_i} \Big\{ &-\log p(s_i[N,\ldots,T \mid \mathbf{x}, s_i[0,\ldots,N-1], \mathbf{s}_{\{i\}^C}[0,\ldots,T]) \\
&+ \gamma_1 |\mathbf{m}_i|_1 + \frac{\gamma_2}{2} |\mathbf{m}_i - \mathbf{m}_i'|_2^2 + \gamma_3 \sum_j |\tilde{\mathbf{c}}_i^{(j)}|_2 \Big\},
\end{aligned}
$$

and

$$
\arg \min_{\tilde{\mathbf{h}}_i, \tilde{\mathbf{f}}_i, \tilde{\mathbf{c}}_i^{(j)}, b_i} \Big\{ -\log p(s_i[N,\ldots,T \mid \mathbf{x}, s_i[0,\ldots,N-1], \mathbf{s}_{\{i\}^C}[0,\ldots,T]) + \gamma_3 \sum_j |\tilde{\mathbf{c}}_i^{(j)}|_2 \Big\}.
$$

Each of the problems were solved using FISTA [26] with the proximal operator for the $L_{2,1}$ penalty from [29].

##### A.5.4 GLM time basis functions

Proper selection of temporal basis functions for the stimulus time filter, spiking feedback filter, and coupling filters is necessary for well-fitted GLM models. As in [10], basis functions corresponding to the first lobe of

$$
b^{(l)}[t] = \frac{1}{2} \cos(a \log[t+c] - \phi_l) + \frac{1}{2} \tag{13}
$$

were used. The quantities, $a$, $c$, and $\phi_l$, as well as the number of basis functions used are hyperparameters. Because the GLM was fit with a 1 ms bin width and the filters were 250 samples long, $t$ takes values $0, 1, \ldots, 249$. The paper used the following basis hyperparameters:

- For the stimulus time filter, $a = 5.5$, $c = 1.0$, and 10 total basis functions were used, corresponding to $\phi_l \in \{\frac{8\pi}{2}, \frac{9\pi}{2}, ..., \frac{16\pi}{2}, \frac{17\pi}{2}\}$.
- For the feedback filter, $a = 5.5$, $c = 1.0$, and 18 total basis functions were used, corresponding to $\phi_l \in \{0, \frac{\pi}{2}, ..., \frac{16\pi}{2}, \frac{17\pi}{2}\}$.
- For the coupling filter, $a = 3.2$, $c = 1.0$, and 10 total basis functions were used, corresponding to $\phi_l \in \{0, \frac{\pi}{2}, ..., \frac{9\pi}{2}\}$.

The basis set for the stimulus time filter was uniformly shifted backward in time to approximately match each retina's response latency to the stimulus.

###### A.5.5 Derivation of the total encoding log-likelihood term (equation (6))

The log-likelihood of the spike trains of all of the RGCs given the stimulus image and the spikes occurring prior to presentation of the stimulus can be computed using the chain rule. In particular, the total log-likelihood over all of the cells corresponding to time bin $t$

$$\log p(\mathbf{s}[t] \mid \mathbf{x}, \mathbf{s}[0, \dots, t-1]) = \sum_{i \in \text{cells}} \log p(s_i[t] \mid \mathbf{x}, s_i[0, \dots, t-1], \mathbf{s}_{\{i\}^C}[0, \dots, t-1])$$

can be computed simply by summing over the cells, since the spiking responses in time bin $t$ for each cell are independent of the other cells given the image and the spike histories for every cell up until time $t-1$. The overall log-likelihood corresponding to time bins $N, N+1, ..., T$ can then be computed using the chain rule

$$\begin{aligned}\log p(\mathbf{s}[N, ..., T] \mid \mathbf{x}, \mathbf{s}[0, ..., N-1]) &= \sum_{t=N}^{T} \log p(\mathbf{s}[t] \mid \mathbf{x}, \mathbf{s}[0, ..., t-1]) \\ &= \sum_{t=N}^{T} \sum_{i \in \text{cells}} \log p(s_i[t] \mid \mathbf{x}, s_i[0, ..., t-1], \mathbf{s}_{\{i\}^C}[0, ..., t-1])\end{aligned}$$

##### A.6 Initialization of HQS for MAP-GLM-dCNN method

We tested different methods of initializing $\mathbf{z}^{(1)}$ in Algorithm 1 for the MAP-GLM-dCNN. We compared initialization with the linear solution (as used for all results shown in the main text), with Gaussian noise initialization with standard deviations $\sigma \in \{10^{-4}, 10^{-3}, 10^{-2}, 10^{-1}, 1.0\}$, where images are defined on the interval $[-1, 1]$. Example test reconstructions for each initialization are shown in Figure 4, and mean test MS-SSIM and PSNR values for each initialization method are summarized in Tables 6 and 7. Qualitatively, the content and structure of the reconstructions did not show significant dependence on the initialization (with exception of the horizontal antenna of the insect in row C). However, the mean luminance of the image reconstructions did depend on the initialization method. Specifically, initialization with the linear solution tended to better recover the mean luminance (rows C, J, N) in cases where the stimulus image was particularly bright or particularly dark. Quantitatively, mean test and heldout MS-SSIM was largely independent of the initialization method (Table 6), while PSNR was worse for the randomly-initialized reconstructions (Table 7).

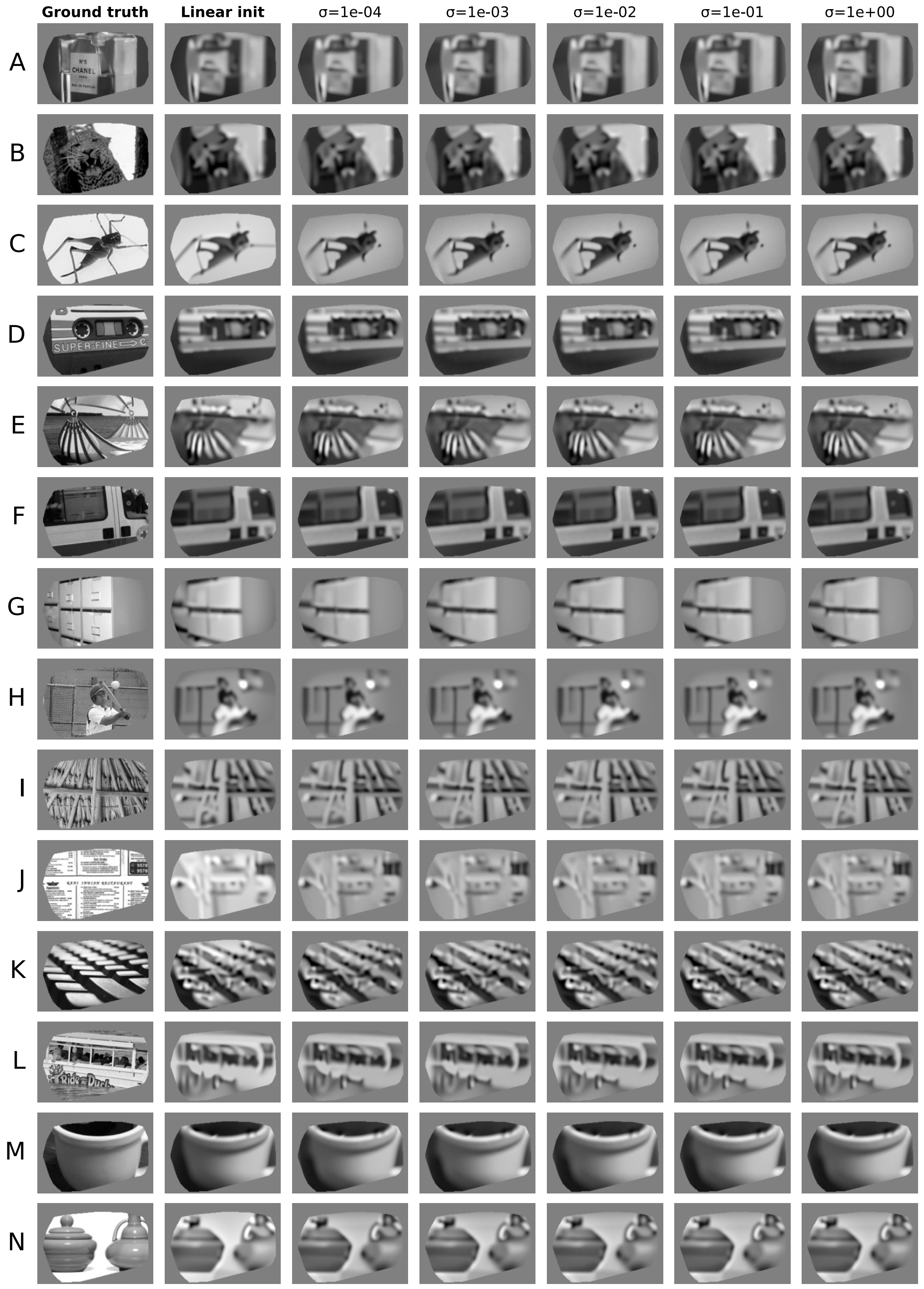


Figure 4: Example MAP-GLM-dCNN reconstructed images for the different initialization methods for $\mathbf{z}^{(1)}$ in Algorithm 1. Column contents: (1) Ground truth stimulus image; (2) Initialization with the linear solution, the method used in the main text; (3) random Gaussian initialization with $\sigma = 10^{-4}$; (4) random Gaussian initialization with $\sigma = 10^{-3}$; (5) random Gaussian initialization with standard deviation $\sigma = 10^{-2}$; (6) random Gaussian initialization with $\sigma = 10^{-1}$;, (7) random Gaussian initialization with $\sigma = 1$. All of the $\sigma$-values are defined for images that lie within the interval $[-1, 1]$.

Table 6: Average test and heldout MS-SSIM for each initialization method, for MAP-GLM-dCNN reconstruction. Initialization with the linear solution, the method presented in the main text, is bolded.

| | **Linear sol.** | | $\sigma = 10^{-4}$ | | $\sigma = 10^{-3}$ | | $\sigma = 10^{-2}$ | | $\sigma = 10^{-1}$ | | $\sigma = 1$ | |
|---|---|---|---|---|---|---|---|---|---|---|---|---|
| | test | held | test | held | test | held | test | held | test | held | test | held |
| Retina 1 | **0.689** | **0.688** | 0.685 | 0.680 | 0.685 | 0.680 | 0.685 | 0.680 | 0.685 | 0.680 | 0.685 | 0.680 |
| Retina 2 | **0.668** | **0.673** | 0.666 | 0.670 | 0.666 | 0.670 | 0.666 | 0.670 | 0.666 | 0.670 | 0.666 | 0.670 |

Table 7: Average test and heldout PSNR for each initialization method, for MAP-GLM-dCNN reconstruction. Initialization with the linear solution, the method presented in the main text, is bolded.

| | **Linear sol.** | | $\sigma = 10^{-4}$ | | $\sigma = 10^{-3}$ | | $\sigma = 10^{-2}$ | | $\sigma = 10^{-1}$ | | $\sigma = 1$ | |
|---|---|---|---|---|---|---|---|---|---|---|---|---|
| | test | held | test | held | test | held | test | held | test | held | test | held |
| Retina 1 | **19.5** | **19.6** | 18.8 | 18.9 | 18.8 | 18.9 | 18.8 | 18.9 | 18.8 | 18.9 | 18.8 | 18.9 |
| Retina 2 | **18.5** | **18.5** | 18.3 | 18.3 | 18.3 | 18.3 | 18.3 | 18.3 | 18.3 | 18.3 | 18.3 | 18.3 |

##### A.7 Implementation details for L-CAE and performance on experimental data

We used the published L-CAE model architecture [8]. This consisted of a linear regression decoder with parameters fitted by solving the normal equations, followed by a 4-layer convolutional encoder and 4-layer convolutional decoder to improve the linear reconstructions. The convolutional encoder and decoder were trained with backpropagation using a masked MSE loss (including only the regions of the image covered by recorded RGCs), the Adam optimizer [53] with learning rate $4 \cdot 10^{-3}$, and batch size 32. The number of training epochs was determined by evaluating the masked MSE loss on the test partition of the dataset.

The results in [8] were based on simulated RGC responses, and to our knowledge there are no published examples applying the L-CAE technique to experimentally recorded data. To verify that the L-CAE performs well with experimental data, we compared the performance of L-CAE and linear regression on the test partition (Figure 5). As expected, the L-CAE systematically produces improved PSNR, relative to linear regression. It also produces more perceptually accurate image reconstructions.

##### A.8 Implementation details for Kim *et al.* linear/nonlinear regression benchmark

The Kim *et al.* paper [4] reconstructs images from RGC spikes by breaking up the problem into three sequential *ad hoc* steps. The target image is decomposed into low spatial frequency and high spatial

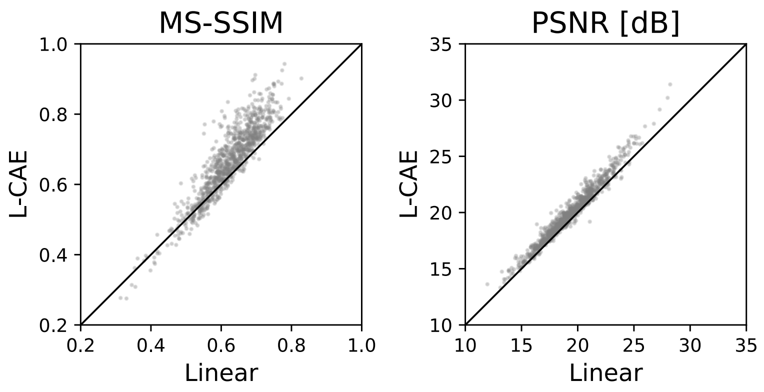


Figure 5: Performance of the L-CAE vs. linear reconstruction on the test partition of the experimental dataset for retina 1, demonstrating that the L-CAE technique can be applied effectively to experimental data. The L-CAE produced systematically greater MS-SSIM than linear reconstruction, suggesting that the L-CAE improves the perceptual similarity to ground truth of the reconstructions. The L-CAE also produced systematically better PSNR than linear reconstruction, which is expected since the L-CAE was trained with an MSE loss.

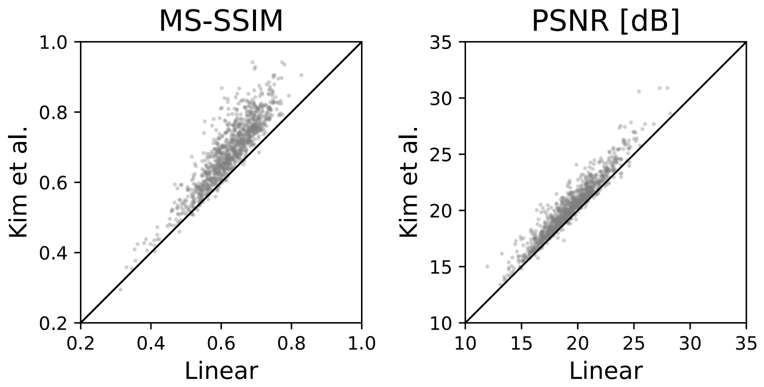


Figure 6: Performance of the Kim *et al.* method vs. linear reconstruction on the test partition of the experimental dataset for retina 1, validating our re-implementation.

frequency components. The low-frequency component is reconstructed using learned reconstruction filters, while the high-frequency component is reconstructed using a trained fully-connected neural network mapping the responses of the most relevant RGCs to the desired high-frequency pixel values. Finally, the low-frequency and high-frequency components are summed together and passed through a deblurring network with an architecture based on that of DeblurGANv2 [54].

The linear reconstruction filter was fitted by solving an L1-regularized linear-least-squares problem aiming to reconstruct the low-frequency image component, where the L1 hyperparameter was chosen by performing a grid search and evaluating MSE on the test partition. Cell/variable selection for the pixel-wise high-frequency component neural network was done by performing a separate L1-regularized linear-least-squares problem aiming to reconstruct the *full* image, with the L1 hyperparameter again chosen using grid search on the test partition. As in the published paper, the top 25 RGCs for each pixel were selected as inputs to the pixel-wise high-pass neural networks by taking the cells with the largest absolute reconstruction filter value for each respective pixel. The high-frequency neural networks were trained with using backpropagation with an MSE loss with respect to the high-frequency component of the target image. The DeblurGANv2 final deblur network was trained using backpropagation with an L1 loss with respect to the target image as well as an L1 perceptual similarity loss computed using the VGG [55] network.

To verify our implementation of the Kim *et al.* method, we compared reconstruction quality for images produced using this method against linear regression (Figure 6). As expected, Kim *et al.* systematically produces better MS-SSIM and PSNR than linear regression.
